# Rapid and accurate protein structure database search using inverse folding model and contrastive learning

**DOI:** 10.1101/2025.05.15.654382

**Authors:** Qiuyi Lyu, Hong Wei, Shuaishuai Chen, Zhenling Peng, Jianyi Yang

**Affiliations:** MOE Frontiers Science Center for Nonlinear Expectations, Research Center for Mathematics and Interdisciplinary Sciences, Shandong University, Qingdao 266237, China; Department of Bioinformatics, School of Basic Medical Sciences, Tianjin Medical University, Tianjin 300070, China; School of Information Science and Engineering, Shandong University, Qingdao 266237, China

## Abstract

Protein structure database search has become increasingly challenging due to the growing number of experimental and computational structures. We introduce mTM-align2, a novel two-step approach for rapid and accurate protein structure database search. In the first step, protein structures are first transformed into embeddings using a pre-trained inverse folding model (ESM-IF) and 3D Zernike polynomials. The ESM-IF embeddings are further optimized through a contrastive learning network, which is trained on ∼7 million structure pairs. Structures with similar embeddings are returned on the fly in this step. The second step employs a rapid structure alignment program to refine top candidates, ensuring high precision and producing high-quality alignments. Extensive benchmarks reveal that mTM-align2 performs competitively compared to other leading methods, completing monomeric structure search in seconds with over 90% precision for the top 10 hits. The t-SNE visualization of the mTM-align2 embeddings for thousands of structures demonstrates that our embeddings are structurally informed, capturing the global structural features. It uncovers insights such as structure misclassifications and ambiguous structural class boundaries. A web server for mTM-align2 is accessible at https://yanglab.qd.sdu.edu.cn/mTM-align/.

## INTRODUCTION

The primary objective of protein structure database search is to efficiently identify similar structures within a structure database, such as the Protein Data Bank (PDB)^1^. The most accurate approach is based on pairwise structure alignment using tools like TM-align^2^ and US-align^3^. However, performing database-wide structure alignments is extremely time-consuming, especially when the database is large. For example, TM-align will require a few weeks to search the PDB database with a single query^4^.

With advancements in protein structure prediction, over 200 million structures predicted by AlphaFold^5^ are accessible in the AlphaFold DB (AFDB)^6^. Searching against these structures is a great challenge. A straightforward approach to accelerate the structure search is to cluster proteins with similar structures. Dali^7^ is one of the most well-recognized methods in this area, which applies a walking strategy to expand hits in clustered structures. mTM-align^8,9^ combines both sequence and structure similarity in the clustering and provides fast pairwise and multiple structure alignment. TM-search^10^ applies an iterative clustering method to reduce the structure comparison and uses TM-align to perform pairwise structure alignment. Although these methods could significantly speed up the search, it is still challenging for them to handle large databases such as AFDB.

To achieve high-speed search, a few methods were proposed using shape-based descriptors ^11-14^, which however are less accurate. 3D-SURFER^14^ compares the global shape similarity using the 3D Zernike Moment^15^. 3D-AF-Surf^11^ improves the performance through deep learning and supports structure search against AFDB. Nevertheless, like 3D-SURFER, it does not consider the residue-level similarity. BioZernike^12^ further enhances retrieval performance through two key innovations. First, it incorporates residue-level information into the moment computation, improving structural representation. Second, it introduces an “alignment descriptor” to facilitate the alignment, leading to a higher precision. In addition to the Zernike moment-based methods, Omakage^13^ employs the incremental distance rank profile to represent proteins shape and the Gaussian mixture model to align protein structures.

Recent works show that deep neural networks have great potential in protein structure representation^16-25^. GraSR^24^ and Progres^23^ utilize a graph neural network to acquire the protein structure embedding, which is further optimized under a contrastive learning framework. DeepFold^18^ and FoldExplorer^20^ use the convolutional/graph attention neural network to encode protein structure, together with sequence information from protein language model. AlphaFind^25^ utilizes the learned metric index^26^ approach to generate protein structure embedding, an extremely compressed representation of protein structure, supporting fast structure search against AFDB. Foldclass^21^ acquires protein structure similarity by comparing their constituent domains, using the program Merizo-search^21^ for domain segmentation. TM-VEC^16^, PLMsearch^17^ and DHR^22^ are representative methods that infer structure similarity from amino acid sequence using protein language models without exact structure comparisons.

Foldseek^4^ is a widely used method for searching various structure databases. It converts each protein structure into an artificial amino acid sequence using a Variational Autoencoder (VAE) network, followed by the use of the MMseqs2^27^ program for quick detection of similar sequences. It was recently extended to search and align multimeric structures, resulting in the new method Foldseek-Multimer^28^.

In this study, we present mTM-align2, an enhanced version of mTM-align tailored for rapid protein structure database search. We utilize the inverse folding model (ESM-IF ^29^) and 3D Zernike polynomials ^12^ in conjunction with a contrastive learning network to improve the speed and accuracy. Benchmarks demonstrate that mTM-align2 is competitive with other methods for searching both monomeric and multimeric structures. The t-SNE visualization shows that the mTM-align2 embeddings effectively capture the global features of protein structures, yielding interesting results, such as identification of potential misclassification of structural classes.

## RESULTS

### Overview of mTM-align2

The major steps of structure search by mTM-align2 is shown in Figure 1. For monomeric structure, the inverse folding model ESM-IF^29^ is used to generate a 2D embedding (512×*L*, where *L* is the length of the structure). This embedding is reduced to a raw embedding (512-D vector) by row-wise sum pooling. The raw embedding from the inverse folding space is then transformed into an updated embedding in the Euclidian space using contrastive learning network (Siamese^30^, see Figure 1c). The similarity between two structures can be calculated on the fly with the cosine function. The top 1000 candidate structures are further filtered and aligned using the fast structure alignment program fTM-align^2^.

**Figure 1.**
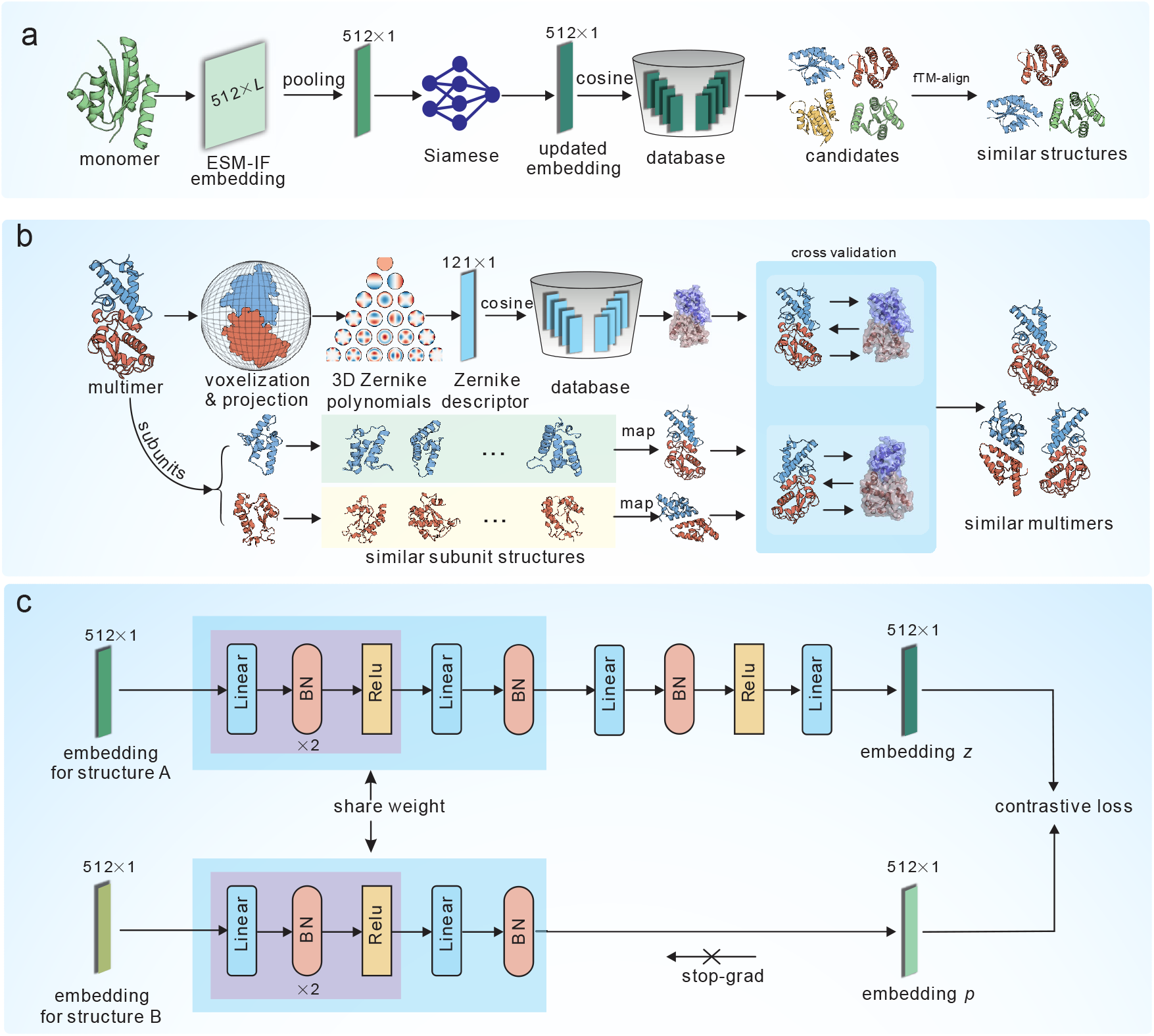
Overall architecture of mTM-align2 for protein structure database search. (a) algorithm for monomeric structure search. The search process begins with the conversion of the query structure into a 2D embedding using the pre-trained inverse folding model (ESM-IF). This 2D embedding is then reduced to a 1D embedding through sum pooling. A contrastive learning network, specifically an asymmetric Siamese network (illustrated in Figure c) is used to optimize the 1D embedding. The similarity between the query and database embeddings is calculated on the fly with cosine function. Finally, the candidates with high similarity to the query are filtered using the structure alignment program fTM-align. (b) algorithm for multimeric structure search (see Figure S7 for more details). It comprises two key modules. The first module utilizes the 3D Zernike polynomials (ZPM) to convert each structure into a 121-D descriptor vector. The similarity between query and database vectors is also calculated by cosine function. The second module is based on monomeric structure search. The monomeric hits for each subunit structure are mapped to their corresponding multimeric structures. The hits from both modules are combined to generate the final set of multimeric structures. (c) Siamese network with an asymmetric architecture to optimize the ESM-IF embedding. For training, the raw embeddings for two structures are fed into the network twice by exchanging their order. For inference, the embedding *z* from the upper branch serves as the final embeddings for structures.

For multimeric structures, two modules are utilized to perform the search (Figure 1b). The first module employs the 3D Zernike polynomials (ZPM)^12^ to identify structures with similar shapes. Each multimeric structure is represented as a 121-D vector derived from ZPM (refer to Methods). The similarity between two vectors is calculated using the cosine function. A maximum of 1000 hits are returned from this module. The second module is based on the monomer search procedure. All subunit structures are first extracted from the multimeric structure. Each subunit is then processed through the monomeric structure search pipeline. The returned monomers are then mapped back to their corresponding multimers, resulting in a maximum of 1000 multimeric hits. Finally, hits from both modules are combined, resulting in the final set of multimers (see Methods).

### mTM-align2 outperforms other methods for multimeric structure search

Here, we compare mTM-align2 with other multimeric structure search methods, including BioZernike^12^, and Foldseek-MM-TM^28^. Foldseek-MM-TM is a variant version of Foldseek-MM^28^ that filters hits using TM-align. We also evaluate the performance of two key modules of mTM-align2: ZPM and the Inverse Folding-based Module (denoted by IFM). The ground truth is defined according to US-align, where a hit is considered a true positive if the TM-score exceeds 0.65, a threshold employed by Foldseek-MM.

The comparison involves searching a set of 286 multimeric structures against a non-redundant database of approximately 30,000 multimeric structures. As shown in Figure 2a, mTM-align2 outperforms Foldseek-MM-TM in terms of precision. For the top 10 to top 50 hits, mTM-align2 achieves precision rates ranging from 55.52% to 35.08%, compared to 48.67% to 26.17% for Foldseek-MM-TM (see Figure S1). The low precision for both methods (< 60%) is likely due to the strict definition of true positives (TM-score > 0.65). When the cutoff is lowered to 0.5, the precisions for the top 10 hits significantly rise to 97.14% and 95.88% for mTM-align2 and Foldseek-MM-TM, respectively.

**Figure 2.**
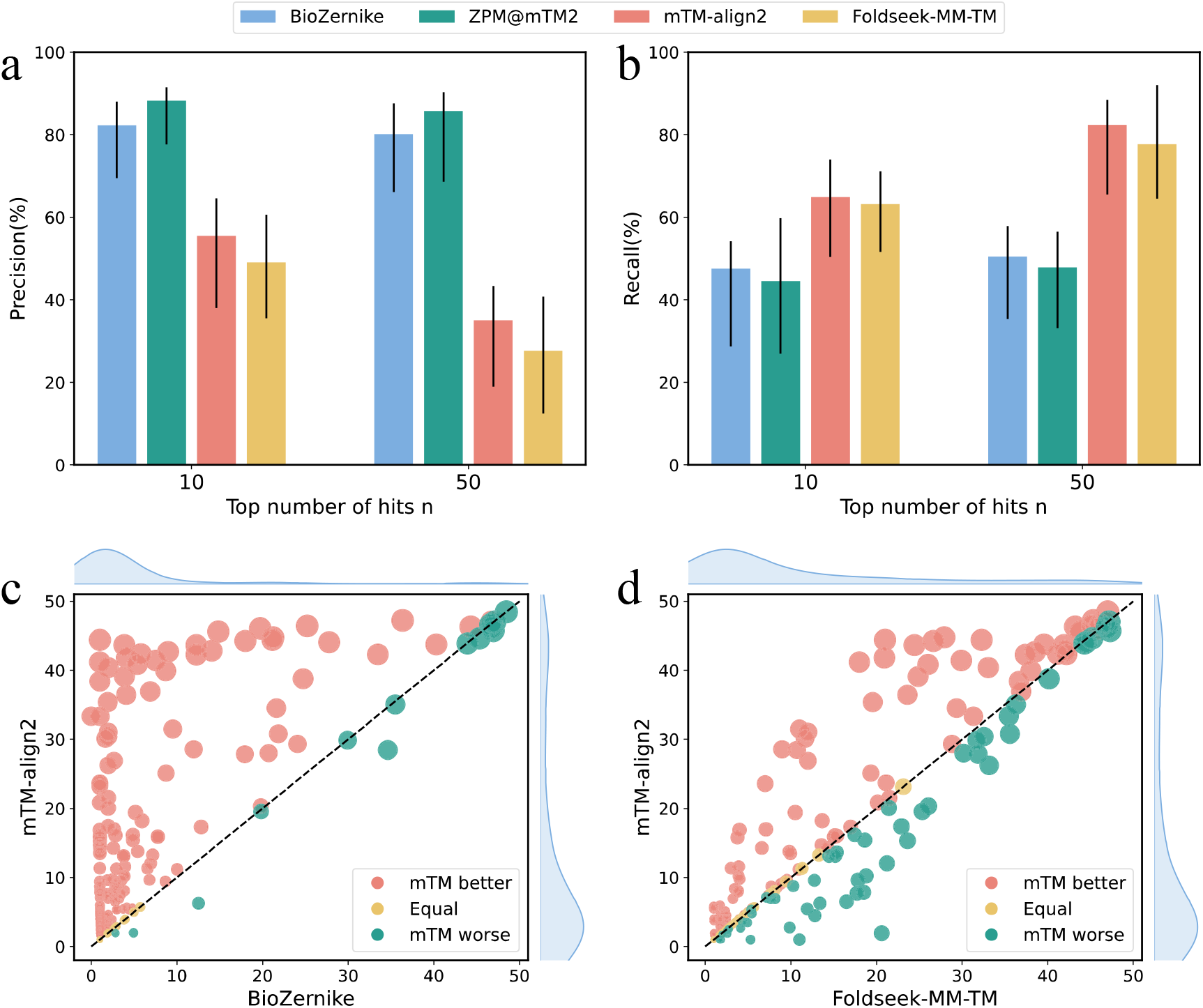
Performance on multimeric structure search. (a) Precision and (b) Recall metrics were evaluated for searching 286 structures in the multimer test set, focusing on the top 10 and 50 hits. ZPM@mTM represents mTM-align2 with ZPM only. (c) and (d) present the summed TM-scores for the true positives among the top 50 hits. Each point in these graphs represents an individual structure from the test set. The radius of each point is proportional to the number of structures retrieved by US-align. Blue lines along the axes are the kernel density estimation of the data.

The Zernike polynomials-based methods (ZPM and BioZernike) demonstrate higher precision than mTM-align2 and Foldseek-MM-TM, at the expense of lower recall. This is because these approaches focus solely on global similarity, often overlooking local structural details and returning a limited number of hits. For instance, the average number of hits returned by ZPM and BioZernike is 13 and 15, respectively, leading to lower recall values (see Figure 2b). Two examples are given to illustrate the limitations of relying solely on Zernike polynomials-based descriptors. In the first example, though the two structures (PDB IDs: 1BAR, 2IOY, Figure S2a) have similar shape (Zernike score 0.96), their local structures are very different with a low TM-score of 0.25. On the contrary, though the two structures (PDB IDs: 8DMG, 5DYP, Figure S2b) have different overall shapes (Zernike score 0.94), they show a high structure similarity (an average TM-score of 0.75). These examples underscore the necessity of integrating Zernike polynomials with other complementary descriptors, such as those derived from the inverse folding model.

The recall values for all methods are summarized in Figure 2b. It needs to be noted that we only consider top 10 to 50 hits in this experiment, thus, the average recall rate may be low. To show the upper limit of the recall rate, we use US-align to define the ground truth and present its top 10 to 50 hits recall rate in Figure S1b. Notably, mTM-align2 achieves higher recall than other methods. For the top 10 hits, mTM-align2 secures a recall rate of 64.91%, compared to 63.56% for Foldseek-MM-TM. This difference becomes more pronounced for the top 50 hits, where mTM-align2 achieves a recall of 82.4% versus 77.59% for Foldseek-MM-TM.

We also compare mTM-align2 with BioZernike and Foldseek-MM-TM based on the sum TM-score (denoted by sTM-score) of the true positives among the top 50 hits. Out of the 286 queries, mTM-align2 outperforms BioZernike for 181 queries and performs worse for 14 queries (see Figure 2c). Notably, the improvement over BioZernike is significant for 61 structures, where the difference in sTM-score is more than 10. A substantial number of points (151) are located in the lower left area (sTM-score < 10 for mTM-align2 and BioZernike), indicating that both mTM-align2 and BioZernike return a limited number of true positives due to the novelty of the query structures (released after January 1, 2023) and the database containing a limited number of similar structures. This is supported by the fact that on average only 5 hits are returned by US-align for these targets.

In the comparison with Foldseek-MM-TM based on sTM-score (see Figure 2d), mTM-align2 performs better for 112 structures and worse for 52 structures. Notably, mTM-align2 significantly outperforms Foldseek-MM-TM for 22 structures (with sTM-score difference greater than 10). An example is illustrated in Figure S2c (PDB ID: 8F5F), which is a homo dimer. In this case, 44 structures with TM-scores > 0.65 are identified by US-align. mTM-align2 successfully found 37 of them, compared to only 2 by Foldseek-MM-TM. Two structures that Foldseek-MM-TM failed to recognize (PDB IDs: 4MPN and 4MP7) are shown in the figure. This failure may be attributed to misalignment of certain alpha helices and beta sheets (highlighted in the red box), resulting in different local 3D interaction (3Di) state sequences in Foldseek-MM-TM.

### mTM-align2 is competitive with other methods for monomeric structure search

We compare mTM-align2 with other methods for monomeric structure search, including Foldseek^4^, Foldseek-TM^4^, fTM-align^2^, BioZernik^12^, DALI^7^, MMseqs2^27^. Two variants of mTM-align2, mTM-align2 without fTM-align, and mTM-align2 without the Siamese network, are also assessed here. The comparison is based on a search of 500 structures against a monomeric structure database of approximately 730,000 structures. A hit is defined as a true positive if it shares a TM-score > 0.5 with the query structure, as calculated by TM-align.

The results are summarized in Figure 3. mTM-align2 achieves comparable precision (>95%) to Foldseek-TM (see Figure 3a). This is anticipated because both methods apply a two-step search strategy. The first step is to quickly find candidate structures that are similar to the query; the second step filters the top hits using the accurate but slow structure alignment. When the filtering is removed, both methods experience reduced precision. For example, the precision of the top 10 hits drops from 98.71% to 84.06% for mTM-align2, compared to a reduction from 99.39% to 89.61% for Foldseek. DALI demonstrates slightly better precision and recall compared to mTM-align2_noTM and Foldseek, but at the cost of being orders of magnitude slower.

**Figure 3.**
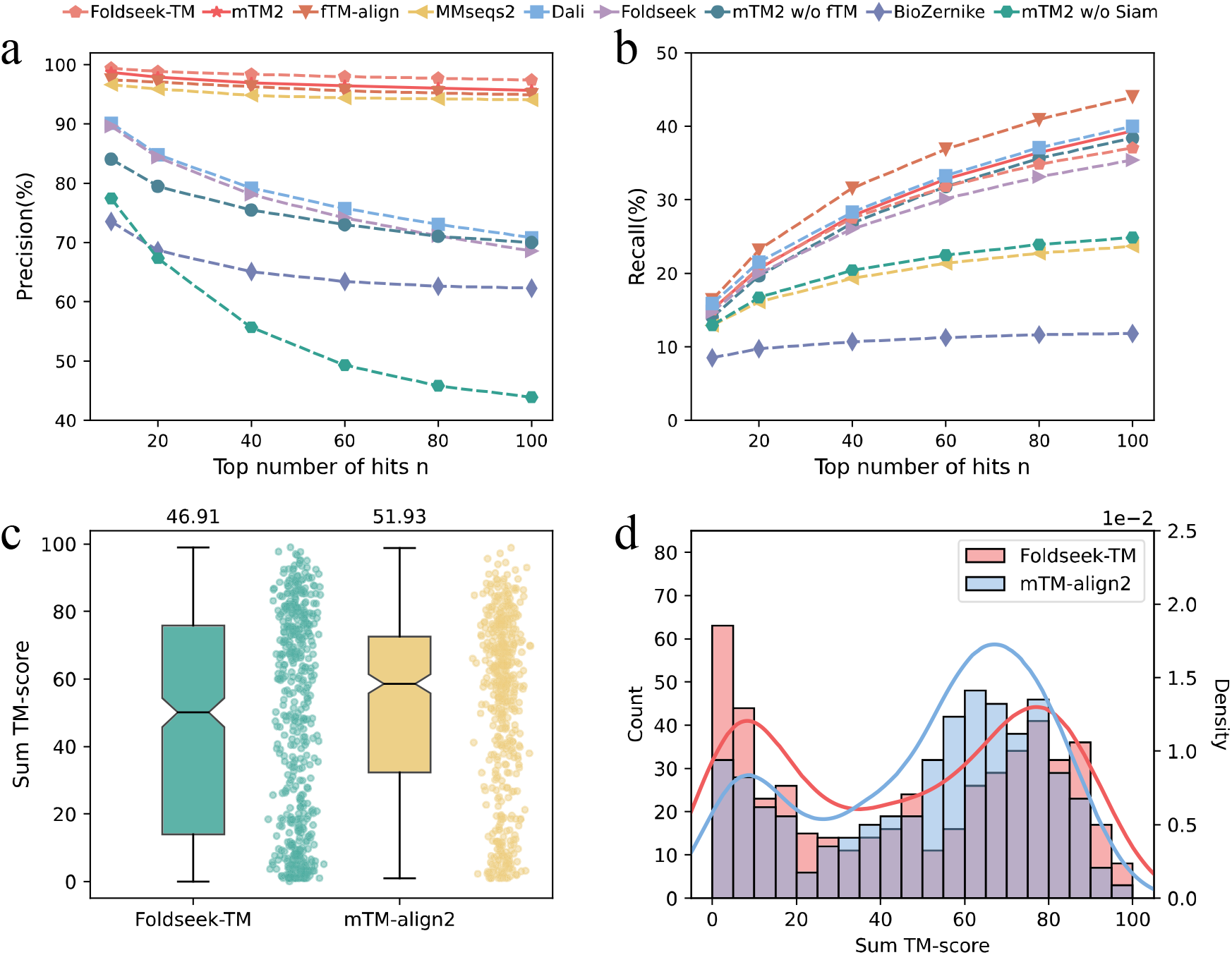
Performance of monomeric structure search. (a) precision and (b) recall metrics were assessed on the monomer test set, which includes 500 structures. (c-d) are the comparisons of the sum TM-scores for the true positives among the top 100 hits. Each point in (c) represents an individual structure from the test set. The average sum TM-scores are listed on the top of box. (d) presents the distributions of the sum TM-score for mTM-align2 and Foldseek-TM, where bars and curves are Count and Density, respectively.

Interestingly, the sequence alignment-based method MMseqs2 has similar precision to mTM-align2 and Foldseek-TM. This may be because it only returns hits with similar sequences, which usually implies similar structures. However, MMseqs2 is not able to detect remote homologies, which share similar structure but dissimilar sequence to the query, as evidenced by its low recall values (see Figure 3b). Note that BioZernike performs poorly in terms of both precision and recall, consistent with previous observations^12^.

We note that the recall values for all methods are low. For instance, the recall value for mTM-align2 is 39.38% even when considering the top 100 hits. To understand this data, we calculate the recall values for the structure alignment-based method fTM-align, which also yields a low recall value of 44.01% for the top 100 hits. The low recall values can be attributed to the limited number of top hits considered (a maximum of 100). The average and median number of structures sharing a TM-score > 0.5 for the 500 testing structures are 585 and 314, respectively, which is significantly larger than 100 (see Figure S3).

The sTM-scores among the top 100 hits returned by mTM-align2 and Foldseek-TM are presented in Figures 3c and 3d. mTM-align2 has a higher average sTM-score than Foldseek-TM (51.93 vs 46.91). The distributions in Figure 3d indicate that mTM-align2 outperforms Foldseek-TM when the sTM-score is less than 70. Specifically, mTM-align2 performs better than Foldseek-TM for 216 out of the 319 targets that the sTM-scores are less than 70. A notable example is shown in Figure S4, which belongs to the *lipocalin-like beta barrel domain* (PDB ID: 8DML, chain B). For this example, among the top 100 hits returned by mTM-align2 and Foldseek-TM, 97 and 20 are true positives, respectively, yielding respective sTM-scores of 60.12 and 13.43. mTM-align2 successfully identifies 77 true positives that are missed by Foldseek-TM. The superimposition of two of these hits against the query structure reveals that the structures in the common core regions (highlighted in red) are highly similar, while other outlier regions differ significantly.

### mTM-align2 is competitive with other methods for SCOP domain classification

The ground truth for the comparison above relies on structural similarity defined by TM-align, which may introduce bias against structure alignment methods. To address this issue, we further compare mTM-align2 with other methods based on the SCOP^31^ domain classification. The dataset comprises 379 structure domains randomly selected from the SCOPe40^31^ database (version 2.08), which contains 15,172 domains with less than 40% pairwise sequence identity. Hits that match the fold/superfamily/family label of the query are regarded as true positives. A precision-recall curve is plotted by adjusting the scoring thresholds of each method (e.g., predicted TM-score in mTM-align2).

Figure 4 summarizes the precision-recall curves for the fold, superfamily and family classification. Similar to the results in the previous section, mTM-align2, Foldseek-TM, and fTM-align perform the best and are indistinguishable to each other. When the structure-based filtering is removed, both mTM-align2 and Foldseek-TM surfer a performance degradation. The decrease is more pronounced at the fold level compared to the superfamily and family levels. For instance, at a recall of 62%, the precision for both mTM-align2 and Foldseek-TM drops from over 80% to about 65% after removing the filter. In contrast, the corresponding decrease for superfamily and family classifications is less than 5%. This difference may be attributed to the high correlation between the TM-score (used in filtering) and the definition of the SCOP folds.

**Figure 4.**
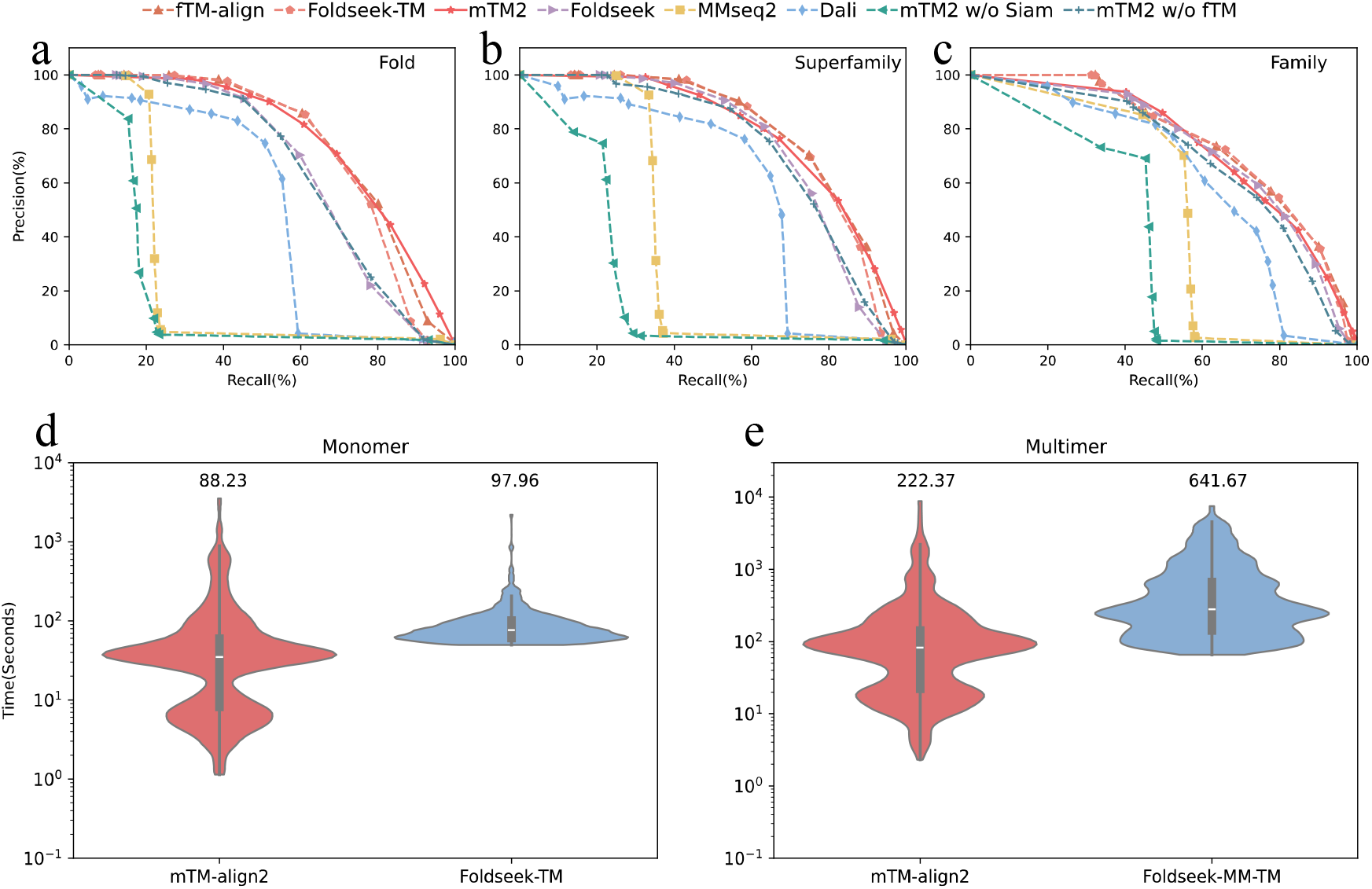
Performance on the SCOPe dataset and running time analysis. (a), (b), and (c) are the precision-recall curves on the SCOPe40 test set, which consists of 379 structures. These comparisons are made at the fold, superfamily, and family labels, respectively. Hits that share the same fold, superfamily, or family labels as the query are defined as true positives. (d-e) compare the average running times for mTM-align2 and Foldseek-TM/Foldseek-MM-TM, evaluated on the monomer and multimer test datasets, respectively.

### mTM-align2 is faster than Foldseek

The running speed of mTM-align2 is compared with the state-of-the-art method Foldseek in Figure 4. Both methods were executed locally on a single core of a Linux server equipped with 4 Intel Xeon Platinum 8260 CPU 96 cores (2.40GHz) and 2TB memory. Note that the default Foldseek-TM and Foldseek-MM-TM utilize all available CPU cores; but for this experiment, we adjusted it to run on a single core for a fair comparison.

The experiment involved searching 286 multimeric structures against ∼300,000 multimers; and 500 monomeric structures against ∼730,000 monomers. Figures 5d and 5e illustrate that mTM-align2 is faster than both Foldseek-TM and Foldseek-MM-TM. The fast speed of mTM-align2 can be attributed to two key factors. First, the similarity is estimated with cosine function in mTM-align2, which is much faster than the *k*-mer based heuristic approach in Foldseek. Second, mTM-align2 utilizes a pre-clustered database, allowing searching against a non-redundant dataset and extending results to other cluster members of the returned hits, similar to the strategy used in mTM-align.

**Figure 5.**
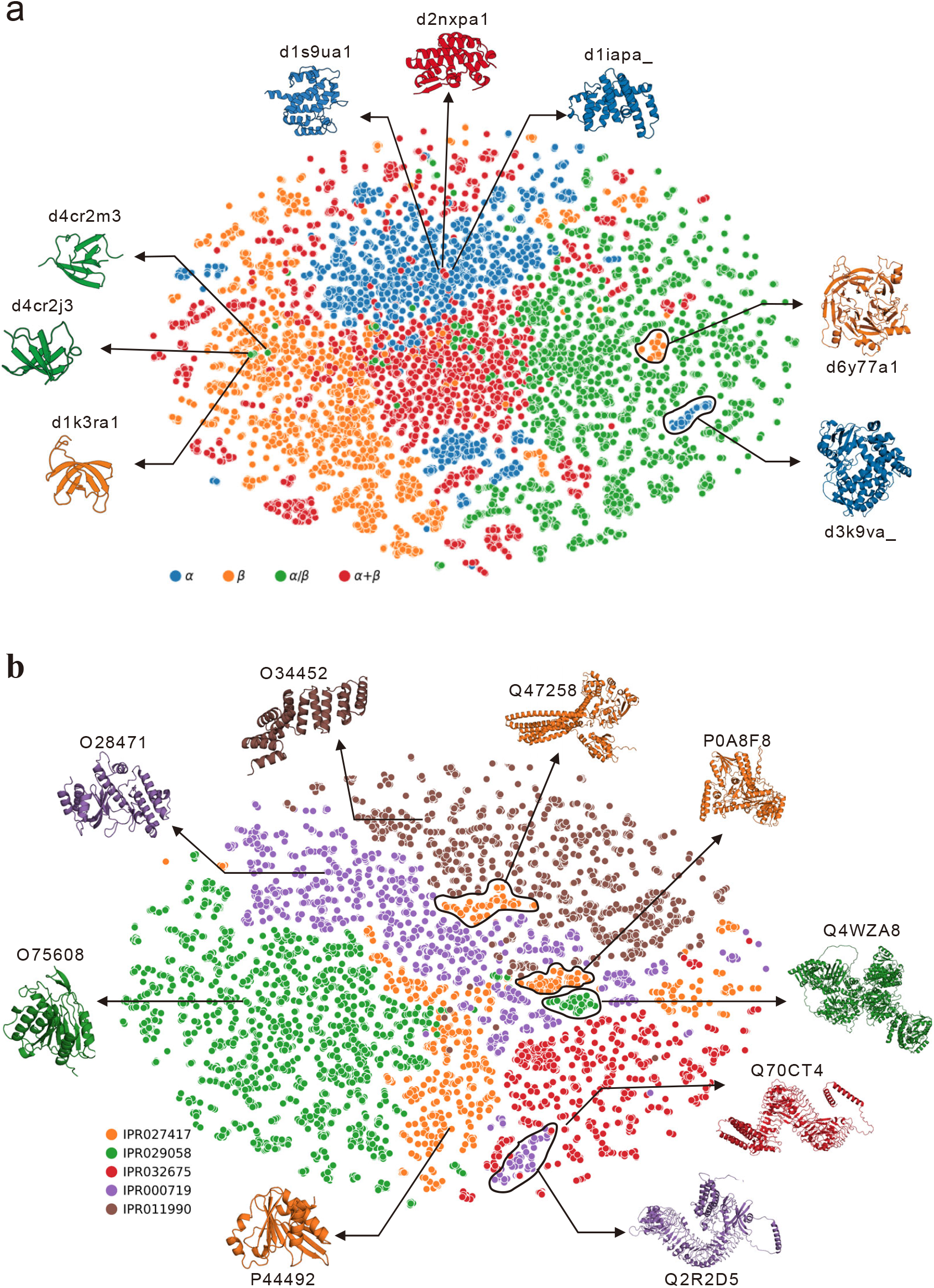
Visualization of mTM-align2 embeddings by t-SNE. (a) results for the 15177 SCOPe structures, where each point represents a structure, colored by structural classes defined by SCOPe. (b) results for 8598 predicted structure models from AFDB. The point colors indicate different InterPro domains.

### t-SNE visualization of the mTM-align2 embedding

Given the outstanding performance of mTM-align2, we visualize its embedding using the t-distributed stochastic neighbor embedding (t-SNE) ^32^ with a perplexity of 5. Figure 5a displays the results for 15177 structures from SCOPe40. Each point in the figure represents one structure, colored according to the structural classes defined by SCOPe. Here, we focus on the top four largest structural classes: α, β, α/β, and α+β. The results indicate that the structures are largely clustered correctly according to their classes, with several noteworthy observations outlined below.

### mTM-align2 embedding reveals misclassification

In the left panel of Figure 5a, two structures (SCOPe IDs: *d4cr2m3* and *d4cr2j3*) belong to the α/β class (green points) but are clustered alongside structures from the β class (orange points). The cartoon visualizations suggest that their structures resemble those of their neighbors, such as the one shown in the figure (SCOPe ID: *d1k3ra1*). These structures primarily consist of β sheets and should be classified as β rather than α/β. This reflects a potential misclassification of structures in the SCOPe database.

### mTM-align2 embedding reveals ambiguous class definition

We also observe that a few α+β-class structures (red points) are clustered with α-class structures (blue points). For instance, the α+β-class structure *d2nxpa1* is grouped with two α-class structures, *d1s9ua1* and *d1iapa_*, as shown at the top of Figure 5a. Although structure *d2nxpa1* contains two small β sheets, its secondary structure is predominantly composed of α helices. This suggests that the class definition for this structure is ambiguous.

### mTM-align2 embedding reveals structural fold

Additionally, we note the emergence of distinct sub-clusters within a single cluster. Two such sub-clusters from the β class (orange points) and the α class (blue points) are highlighted in the right panel of Figure 5a. They appear within the cluster of α/β class structures (green points). Upon inspecting the structures in these two sub-clusters, we present two representative examples (*d6y77a1* and *d3k9va*). Both structures contain α helices and β sheets, indicating shared structural similarities with other α/β-class structures in the cluster. Furthermore, these two sub-clusters correspond to two SCOPe folds, a.104 and b.69, respectively, which have well-defined boundaries with other structures. This suggests that mTM-align2 embedding has the potential to cluster structures from a ‘coarse-grained’ class level to a ‘fine-grained’ fold level.

### mTM-align2 embeddings for predicted structures in AlphaFold DB

We further analyze the predicted structures from AFDB^6^ using the mTM-align2 embeddings. For illustration, we took 8598 high-confidence structures with pLDDT > 80 from five InterPro^33^ domains. Figure 5b demonstrates that most structures cluster well, with clear boundaries between different InterPro domains: IPR029058 (*Alpha/Beta hydrolase*, green points), IPR027417 (*P-loop containing nucleoside triphosphate hydrolase*, orange points), IPR000719 (*Protein kinase*, purple points), IPR011990 (*Leucine-rich repeat domain*, red points), and IPR011990 (*Tetratricopeptide-like helical domain*, brown points). Each InterPro domain is represented by an example structure in the figure, illustrating the structural differences among them.

Notably, the first three InterPro domains (green, orange, and purple points) are closely clustered, likely due to their similar structural features (containing both α helices and β sheets) and functions (enzymes). In contrast, the other two domains (red and brown points) are well-separated, reflecting their distinct structural characteristics: the *Leucine-rich repeat domain* (red points) is characterized by a repeated α/β horseshoe fold, while the *Tetratricopeptide-like helical domain* (brown points) consists of a multi-helical fold made up of two curved layers of α-helices. These results further confirm that the mTM-align2 embeddings are structurally informed.

However, we observe the formation of some sub-clusters, four of which are highlighted in circles. A representative structure is provided for each sub-cluster. A common characteristic of these structures is that they are significantly larger than other structures within their respective domains. For instance, the structure for Q4WZA8 (shown on the right side of Figure 2b) contains over 2,000 amino acids, approximately ten times larger than the example structure O75608 from the same domain. In fact, Q4WZA8 is a multi-domain protein. According to the InterPro annotation, this protein belongs to 21 InterPro domains, including the domain IPR029058 associated with structure O75608. Similarly, the purple structure (Q2R2D5) shown at the bottom of Figure 2b also contains multiple domains, including IPR000719 (purple) and IPR032675 (red). This structure is primarily characterized by the repeated α/β horseshoe fold, leading to its assignment in the cluster of IPR032675 rather than IPR000719. The clustering is further supported by the structural similarity (TM-score > 0.5) between the two representative structures (Q2R2D5 and Q70CT4). These data indicate that the mTM-align2 embeddings effectively capture the global features of protein structures.

### Ablation study

For monomeric structures, mTM-align2 makes use of contrastive learning (Siamese network) and structure alignment-based filter (fTM-align) to enhance the precision. We conducted an ablation study to assess the improvements introduced by these components. On the monomer test dataset, the precision of mTM-align2 without fTM-align drops from 95.64% to 70.11% (Figure 3a, top 100 hits), indicating that the structure-based filter is essential. In addition, the precision significantly declines from 70.11% to 43.90% when the Siamese network is removed. These data suggest the critical role of the Siamese network in enhancing the precision of mTM-align2.

For multimeric structures, the monomeric structure search module (IFM) is combined with the ZPM-based module to enhance the search results (Figure 1b). We evaluated the improvement gained from this combination on the multimer test dataset. When mapping multimeric structures using the top hits from IFM, an average of 32.86 hits are obtained. In contrast, when using the ZPM-based search for multimeric structures alone, an average of 15.6 hits are retrieved. The Venn diagram in Figure S5 demonstrates that the two modules are complementary to each other. Both modules have overlap for 12.39 hits and their unique hits (20.47 and 3.21 for IFM and ZPM, respectively). Therefore, combining hits from both modules results in the most accurate results in mTM-align2.

## CONCLUSION

Protein structure database search has become increasingly important and challenging. Building on advancements in protein engineering and deep learning, we developed mTM-align2, a rapid and accurate approach for protein structure database search and clustering. Unlike other methods, mTM-align2 effectively handles both monomers and multimers within a single framework. Protein structures are converted into embeddings using the inverse folding model and the 3D Zernike polynomials. The embeddings are further optimized through a contrastive learning network trained on ∼7 million structure pairs. The embedding significantly accelerates search speed; while the network enhances the accuracy. Comprehensive benchmarks demonstrate the superior performance of mTM-align2 for both monomers and multimers. mTM-align2 typically completes monomeric structure searches against existing databases within seconds, achieving over 90% precision for the top 10 hits.

The t-SNE visualization of the mTM-align2 embeddings for thousands of protein structures indicates that the mTM-align2 embeddings effectively capture the global features of protein structures, leading to insightful observations, such as identification of potential misclassifications of structural classes and recognition of blurred class definitions. These findings highlight the utility of mTM-align2 in advancing our understanding of protein structures and their classifications and functions.

Despite its strengths, we admit certain limitations of mTM-align2. The sum pooling transform of the ESM-IF embedding can result in the loss of structural details, bringing false positive hits. To solve this problem, we apply structure alignment to the top hits to filter out false positives. However, this filtering slows down the search. Potential solution could involve directly comparing the original ESM-IF embeddings with contrastive learning without pooling. We plan to explore this enhancement in our future work.

## METHODS

### Training and test datasets

#### Monomer test datase

We obtained a dataset of ∼730,000 monomers from Q-BioLiP^34^ (version 2023.01.11) and 15,172 domains from the SCOPe40 database (version 2.08). These structures, comprising both monomers and domains, were clustered using CD-HIT^35^ at a 40% sequence identity threshold to eliminate redundancy, resulting in 70,270 distinct clusters. We randomly selected 500 monomers from 500 clusters as the monomer test set. The test set for the SCOP domain data set was constructed by selecting each domain structures in the same cluster, resulting in 379 domains (some of the selected clusters do not contain any domain structures).

#### Multimer test dataset

We obtained 309,192 multimers from Q-BioLiP (version 2023.01.11). The multimer test set was constructed based on the monomer test set by mapping the selected monomers to their corresponding multimers. From this mapping, 286 multimers are selected to form the multimer test set.

#### Training set

To train the Siamese neural network, we constructed >7 million pairs of protein structures as follows. First, we generated random protein pairs from the non-redundant set of monomeric structures (testing structures were removed), resulting in ∼5 million pairs of structures. Most of them are negative training samples with a TM-score less than 0.5. We then use fTM-align to search for protein pairs with high structure similarity. This step yields a high-similarity set of ∼2 million pairs of structures, each with a TM-score greater than 0.5.

### Algorithm for monomeric structure search

The monomeric structure search involves three key steps (see Figure 1):

#### Step 1. Encode a structure using the inverse folding model

The pre-trained protein language model ESM-IF^29^ is used to transform a monomeric structure into a raw embedding (*L*×512). Note that only the encoder from ESM-IF is used. The embedding of the structure is reduced to a *raw embedding* (512-D vector) by row-wise sum pooling.

#### Step 2. Optimize the structure embedding with contrastive learning

the raw embedding from the inverse folding space is then transformed into an *updated embedding* in the Euclidean space using contrastive learning network (Siamese^30^, Figure 1c, introduced below). As shown in Figure 1c, the Siamese network consists of two branches. For inference, the raw embedding from ESM-IF is fed into the upper branch, producing the updated embedding ***z***.

#### Step 3. Search and filter similar structures

The similarity between two embeddings *z*_1_ and *z*_2_ is defined as the cosine of the angle (*θ*) between ***z***_1_ and ***z***_2_, which is named as IF-score:

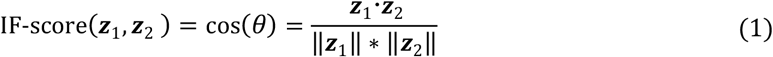

The IF-score between the query embedding and the pre-calculated embeddings of all structures in the database can be calculated on the fly. Structures with IF-score greater than 0.4 are returned. To improve the precision and generate pairwise alignment, we apply the fast protein structure alignment program fTM-align to filter the structures, removing those with a TM-score below 0.4. The retained hits are then ranked and mapped to their respective clusters to expand the search to the whole database. Details for the monomer structure search algorithm can be found in the supplementary algorithm A1.

### Contrastive learning using the Siamese neural network

We use contrastive learning, specifically, the Siamese neural network^30^ with asymmetric structure to optimize the relationship between vector similarity and structure similarity. The network is shown in Figure 1c. During training, as the network is asymmetric, we input the raw features twice by swapping their order. For a structure pair (*x*_1_, *x*_2_), in the first run, we feed the embeddings of *x*_1_ and *x*_2_ to the upper and lower branch, respectively. The order of *x*_1_ and *x*_2_ is swapped in the second run. As a result, we obtain two different vectors for each structure, that is *z*(*x*_1_), *p*(*x*_1_) for *x*_1_, and *z*(*x*_2_), *p*(*x*_2_) for *x*_2_. Then the average cosine similarity (i.e., average of cos(*z*(*x*_1_), *p*(*x*_2_)) and cos(*z*(*x*_2_), *p*(*x*_1_))) is used to estimate the structure similarity (i.e., TM-score) of *x*_1_ and *x*_2_. We show the details of the training process in the supplementary algorithm A2.

*Loss function*. To minimize the differences between our predicted score and the TM-score, the MSE loss is used.

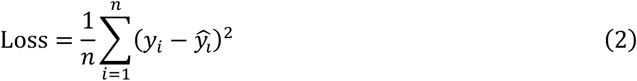

where *y*_i_ is the predicted TM-score, 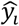 is the real TM-score, *n* is the size of the training batch.

#### Model training

The training and test are conducted on a Linux server equipped with 4 Intel Xeon Platinum 8260 CPU 96 cores (2.40GHz) and 2TB memory. The GPU used for the training was a Nvidia A100 with 40GB of high-bandwidth memory. To achieve better performance, we conduct label-optimization according to the significance of TM-score^36^. For dissimilar protein pairs (TM-score < 0.3), we subtract 0.2 from their TM-score, further separating them in the feature vector space; if the TM-score is above 0.7, we add 0.2 to their TM-score, with a maximum value of 1. This adjustment in the label assignment significantly enhances the retrieval accuracy (Figure S6).

The initial learning rate is 0.001, which is scheduled by a cosine function as the training going on. We utilized a batch size of 1024. Stochastic gradient descent (SGD) was applied to optimize the parameter. The network converged after 100 epochs in about 1.5 hours.

### Algorithm for multimeric structure search

The multimeric structure search consists of two modules. The first module is based on 3D Zernike polynomials (ZPM, introduced below in detail). The second module is based on the monomeric structure search introduced above (denoted by IFM). The hits from both modules are combined to yield the final set of similar multimers. More details are available in the supplementary algorithm A3.

#### Descriptors from 3D Zernike polynomials

We use the 3D Zernike moments^15^ to describe the shape of multimeric structures, with implementation by the package BioZernike^12^. The first step involves representing the shape in 3D space through a process called voxelization. For a given protein, a 3D grid of size 32×32×32 is created to convert its structure coordinates into a volumetric representation. For each amino acid, the Cα atom is used as the representative atom. Then, Gaussian density is constructed for each Cα atom, where the weight corresponds to the amino acid’s molecular weight, and the size reflects the spherically averaged size of the amino acid. Then the Gaussian densities are placed into the volume, which is subsequently scaled to fit within a unit sphere. Finally, a coordinate system is fixed with the origin as the center of the grid and the axes aligned with the grid axes.

After establishing the coordinate system, any given protein can be represented as a volumetric function *f*(***x***). The 3D Zernike polynomials 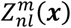 are used as orthonormal basis functions, allowing the volumetric function *f*(***x***) to be decomposed accordingly.

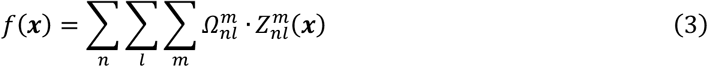

where ***x*** is the vector representing the coordinates of grid points. The coefficients of the basis functions 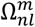 are defined as the 3D Zernike moments, which will be transformed into the shape descriptors after normalization. The 3D Zernike polynomials 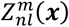 is defined as^15^:

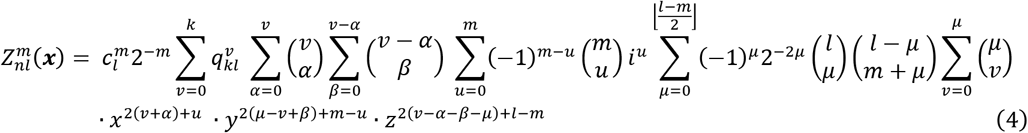

where *i* is the imaginary unit, *n* is the predefined maximum polynomial order, *l* ∈ [0, *n*], *m* ∈ [-*l, l*], *n l* is even number, and 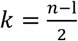. The term 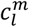 and 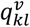 are defined as follows:

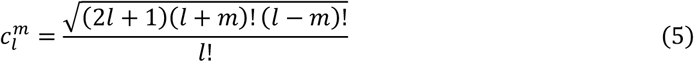

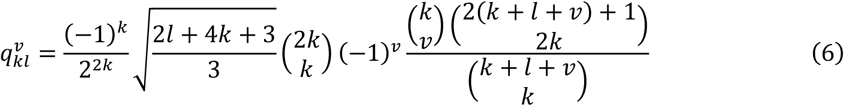

Thus, the 3D Zernike moments are defined based on the volume function *f*(***x***) as

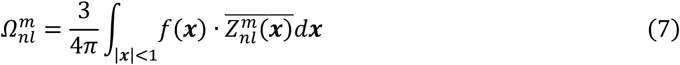

where 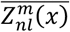 is the conjugate of 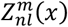

Note that the moment 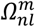 is not invariant under rotation. To obtain rotation invariant shape descriptors, the 3D Zernike descriptors *D*_*nl*_ are defined as the norms of 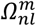, that is

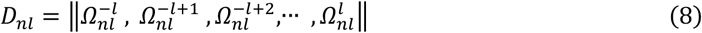

Moments up to order 20 (*n* 20) are utilized in our experiments and the resulting moment is transformed into a 121-D descriptor. This vector describes the overall shape of the structure. The similarity (denoted by ZP-score) between two descriptor vectors is calculated using the cosine function, similar to Equation (1). The structures in the database are ranked by ZP-score and a maximum of 1000 multimeric structures are returned by ZPM.

#### Identify multimeric structures based on monomeric structure search

We make use of the monomeric structure search module to identify multimeric structure (see Figure S7 for more details). The top 6 longest subunit structures are first extracted from the query structure. Each subunit structure is then fed into the monomeric structure search pipeline. The returned monomers are then mapped to their respective multimers, resulting in a maximum of 1000 multimeric hits. The IF-score for each subunit in the mapped multimers is taken from the previous set of similar subunits, which is set to 0 if not existed in the set. The IF-score for each mapped multimer is then calculated as the mean IF-scores over all subunits.

#### Strategy for combining multimeric structure hits from ZPM and IFM

The candidate multimeric structures from ZPM and IFM are combined to generate the final set of multimeric structures based on a consensus score called Q-score (see Figure S7 for more details).

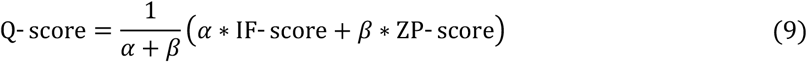

where the weights α and β are empirically set to 1 and 0.3, respectively. Multimeric structures are ranked based on the Q-score and up to 1000 top hits are returned.

### Controlled baselines

We compare mTM-align2 with several existing methods, including fTM-align^2^, a fast implementation of TM-align; DALI^7^, a popular structure alignment-based method using distance matrix; BioZernike^12^, an alignment-free method that is officially used by PDB; Foldseek^4^, the state-of-the-art method for efficient protein structure database search; MMseqs2^27^, an efficient method for fast sequence database search. As the model weight for BioZernike is not publicly available, we manually submitted the query structures to its web server (http://shape.rcsb.org/) to conduct the structure search. For all other methods, we downloaded the packages and ran them locally with default settings.

### Evaluation metrics

Similar to previous studies^9^, the average precision and recall are used to evaluate the performance:

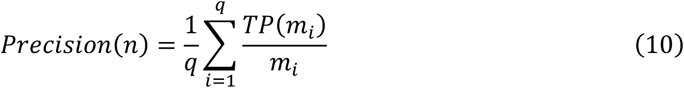

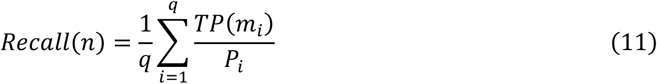

where *q* is the total number of structures in a test set, *n* is the number of top hits to be assessed, *m*_i_ = min(*N*_i_, *n*) and *N*_i_ is the number of returned hits, *P*_i_ is the number of hits that have TM-score > 0.5 with the query. A structure is defined as a true positive (TP) if it appears in the top *n* hits and its TM-score with the query exceeds the specific thresholds. For monomeric/multimeric structures, the threshold is defined as 0.5/0.65. In this study, the TM-score between two structures is defined as the average TM-score, that is, the average of the two TM-scores normalized by the lengths of two structures. Note that TM-align/US-align is used to calculate the TM-score between two monomeric/multimeric structures.

## AVAILABILITY

The web server is available at: https://yanglab.qd.sdu.edu.cn/mTM-align/.

## ACKNOWLEDGEMENT

This work is supported by the National Key Research and Development Program of China (2023YFF1204003), the National Natural Science Foundation of China (NSFC T2225007, T2222012, 32430063), the Shandong Provincial Natural Science Foundation Youth Found (ZR2023QF156), and the Fundamental Research Funds for the Central Universities.

