## supplemental information for "Rapid and accurate protein structure database search using inverse folding model and contrastive learning"

### Supporting Information

#### Supporting Algorithms

##### A1: Algorithm for monomeric structure search

**Algorithm 1** Monomer Retrieval Method for mTM-align2

**Input:** Monomer Coordinate File for *Query*
**Output:** Similar Structure Monomers

```
Residue_q = ESM_IF(Query)
ESM_q = Sum(Residue_q)
//Only feature z from upper branch is used
z_q = Simsiam(ESM_q)
for monomer t in database do
    score_t = cosine(z_q , z_t)
R = monomers that score > 0.4
//Filter the result with TM-align
R_tm = monomers in R with TM-score > 0.4
R_sort = Sort(R_tm)
Result = map monomers in R_sort to its cluster
Return Result
```

##### A2. Details about the Siamese network

**Algorithm 2** Training Method for Siamese Network

**Input:** Momomer Coordinate File for Two Proteins

```
//f: the projector in figure 1c.
//h: the predictor in figure 1c.
for protein pairs (x_1 , x_2) in training batch do
    z_1 , z_2 = f(x_1) , f(x_2)
    p_1 , p_2 = h(z_1) , h(z_2)
    //compute the similarity of descriptors
    s = avg(cos(p_1 , z_2) , cos(p_2 , z_1))
    //compute the loss
    loss = MSE(s , TM-score(x_1 , x_2))
    loss.backward()
procedure cos(p , z)
    //stop gradient
    z = z.detach()
    return cosine(p , z)
```

#### A3: Algorithm for multimeric structure search

**Algorithm 3** Oligomer Retrieval Method for mTM-align2

**Input:** Oligomer Coordinate File for *Query*
**Output:** Similar Structure Oligomers

//Shape Similarity Retrieval
$z_q = Zernike(Query)$ //Zernike Descriptor for Query
**for** *Oligomer o's Zernike Descriptor* $z_o$ *in database* **do**
    $shape(o) = cosine(z_q\,, z_o)$ //compute cosine similarity
//Chain Similarity Retrieval
$chains = \{6\ longest\ chains\ from\ Query\}$
**for** *chain c in chains* **do**
    //retrieve with descriptor from Simsiam
    $Residue_c = ESM\_IF(c)$ //Residue feature for c
    $ESM_c = Sum(Residue_c)$ //Raw embedding for c
    $s_c = Simsiam(ESM_c)$ //Updated embedding for c
    //compute cosine similarity with descriptor from Simsiam
    **for** *chain t's descriptor* $s_t$ *in chain database* **do**
        $simsiam_c(t) = cosine(s_c\,, s_t)$
    //retrieve with Zernike descriptor
    $z_c = Zernike(c)$ //Zernike Descriptor for c
    //compute cosine similarity with chain Zernike descriptor
    **for** *chain t's descriptor* $z_t$ *in chain Zernike database* **do**
        $zernike_c(t) = cosine(z_c\,, z_t)$
    **for** *chain t that* $simsiam_c(t) > 0.60$ *and* $zernike_c(t) > 0.95$ **do**
        $o = map\ t\ to\ its\ oligomer$
        $chain_o(t) = simsiam_c(t)$
//compute the similarity score for oligomers in database
**for** *oligomer o in database and t belong to o* **do**
    $score(o) = (shape(o) + 0.3 * avg(chain_o(t)))\ /\ 1.3$
$result = Sort(score)$ //Sort the result according to score
*Return result*

#### Supporting Figures

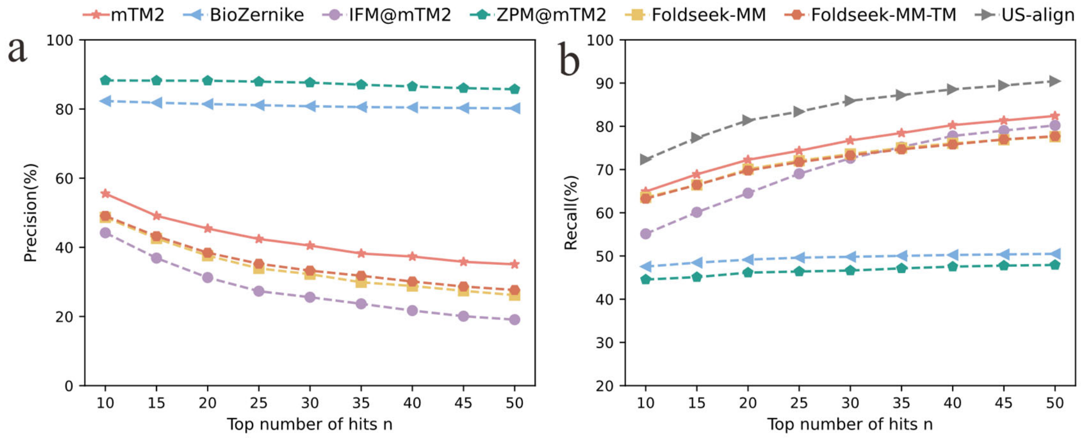


**Figure S1. Performance on multimeric structure search.** (a) Precision and (b) Recall metrics were evaluated for searching 286 structures in the multimer test set, focusing on the top 10 to 50 hits. IFM@mTM and ZPM@mTM represent mTM-align2 with only IFM and ZPM, respectively.

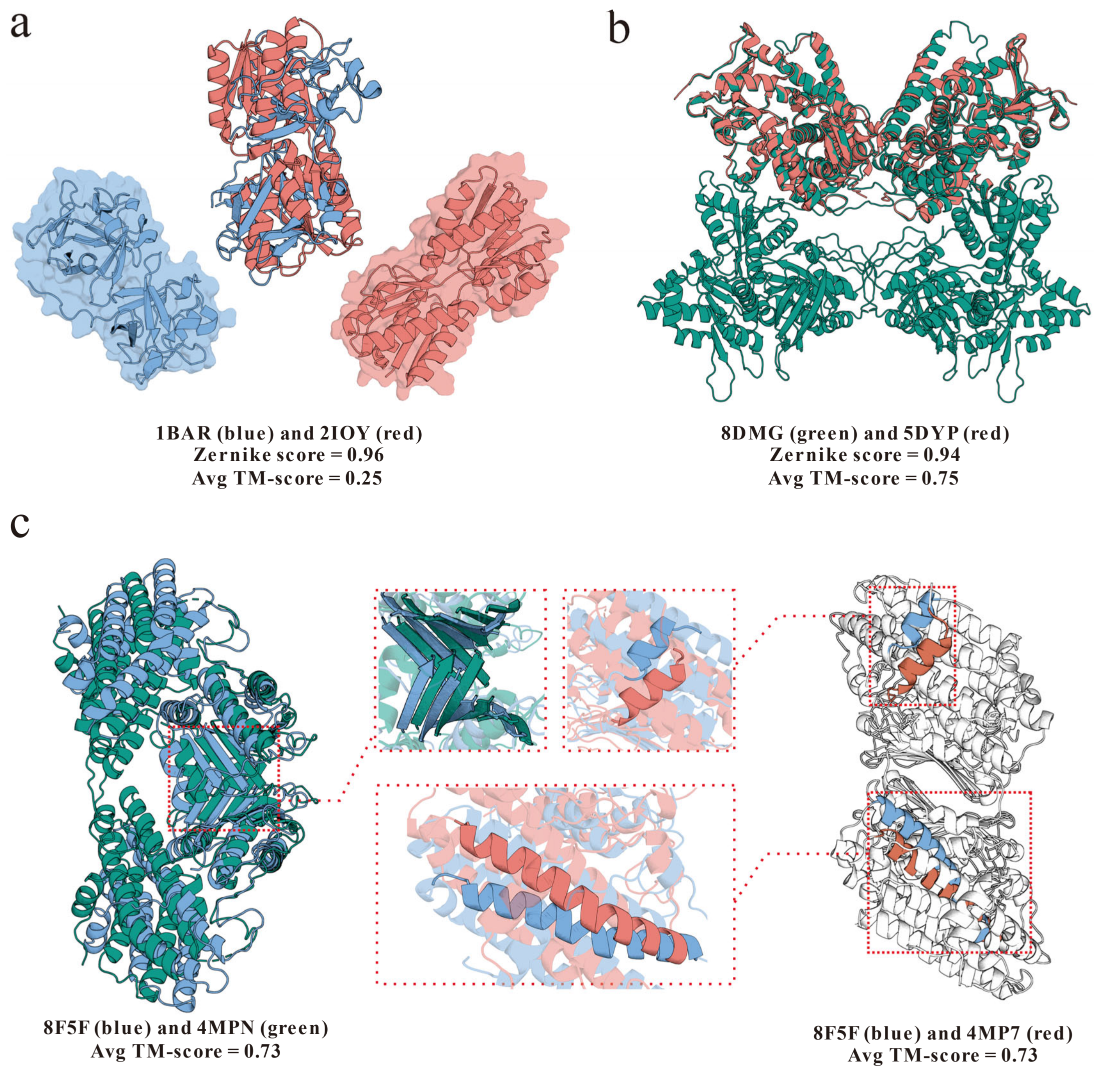


**Figure S2. Case studies for multimeric structure search.** (a) for the query structure (PDB ID: 1BAR, blue cartoon), ZPM returns a false positive hit (PDB ID: 2IOY, red cartoon). The Zernike score between both structures is 0.96 (above the threshold 0.95), while the TM-score is only 0.25 due to very dissimilar structural details. (b) is a false negative example given by ZPM, in which the Zernike score between the query structure (PDB ID: 8DMG, green cartoon) and the returned structure (PDB ID: 5DYP, red cartoon) is 0.94 (blow the threshold 0.95). However, the average TM-score is 0.75, indicate that they share similar structures. (c) shows two false negative examples from Foldseek-MM-TM. The query structure (PDB ID: 8F5F, blue cartoon) share similar structure with two templates: PDB ID: 4MPN, TM-score 0.73; and PDB ID: 4MP7, TM-score 0.73. However, Foldseek-MM-TM failed to detect these structures.

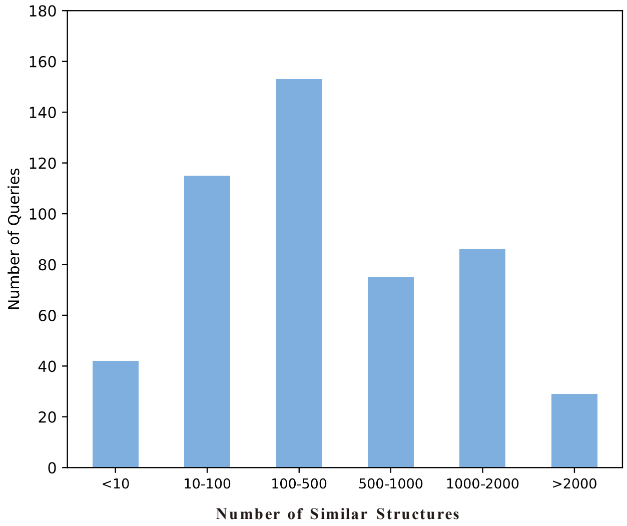


**Figure S3. The number of similar structures (by TM-align) for the 500 monomeric test structures.** Most query structures have more than 100 similar structures in the database. However, up to 100 hits are assessed, resulting in low recall for all methods.

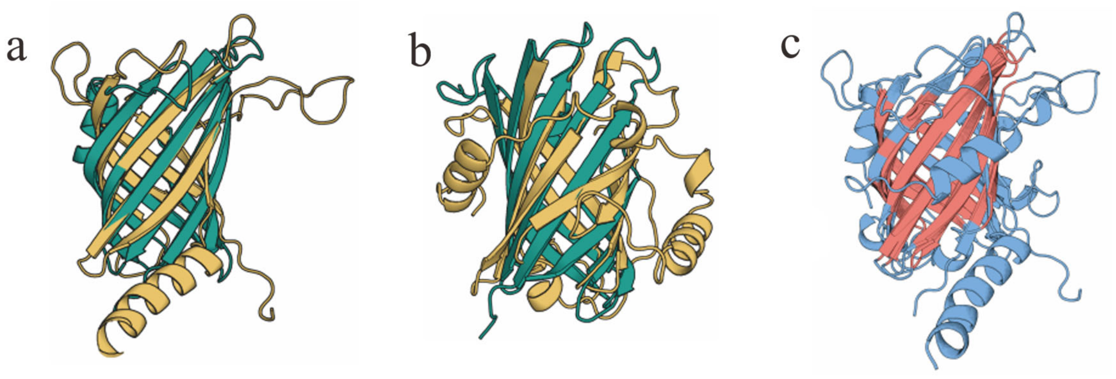


**Figure S4. An example monomeric structure showing that hits missed by Foldseek-TM are successfully detected by mTM-align2.** The query structure, 8DML_B, shares over 0.5 TM-score with two hits 1MM4_A (TM-score: 0.54) and 4GET_D (TM-score: 0.52). They are however missed by Foldseek-TM but successfully identified by mTM-align2. (a) displays the superposition of 8DML_B (green) and 4GET_D (yellow). (b) shows the superposition of 8DML_B (green) and 1MM4_A (yellow). (c) multiple structure alignment for three structures using mTM-align2, where the common core regions are highlighted in red.

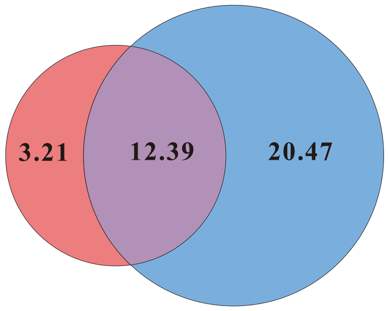


**Figure S5. Venn diagram for the average number of results return by ZPM (red) and IFM (blue) on multimer test dataset.** The number of common results returned by the two module is shown in purple. ZPM focus on global shape similarity. A protein will be returned as a hit only if its ZP-score is larger than 0.95. On the contrary, IFM considers more on residue level structure information. A protein is returned as a hit if its IF-score is larger than 0.6.

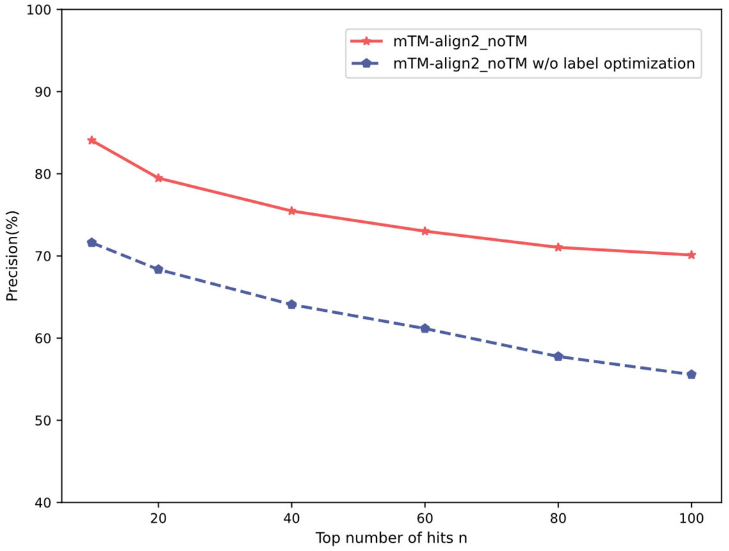


**Figure S6. Ablation study for label optimization.** To achieve better performance, we conduct label-optimization according to the significance of TM-score. For dissimilar protein pairs (TM-score < 0.3), we subtract 0.2 from their TM-score, further separating them in the feature vector space; if the TM-score is above 0.7, we add 0.2 to their TM-score, with a maximum value of 1.

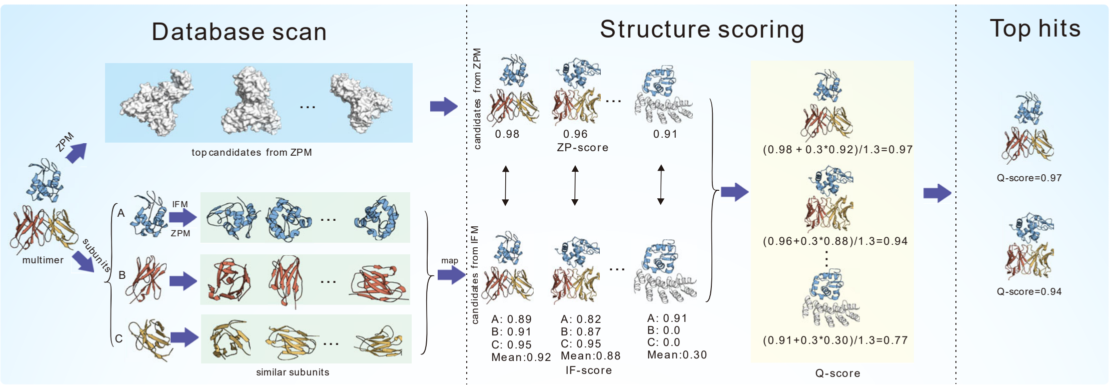


**Figure S7. Flowchart for oligomeric structure search.** Two modules are used here. The first module (ZPM) calculates the shape similarity (ZP-score) between multimeric structures based on 3D Zernike polynomials. Structures with high ZP-scores are returned. The second module splits the input structure into subunits and detects similar monomers using both IFM and ZPM. Specifically, for each subunit extracted from the multimer, we detect similar monomeric structures using the IFM module. There are a few key modifications compared with the monomer search procedure in Figure 1a. We increase the IF-score threshold from 0.4 to 0.6 to enhance precision. In addition, the structure alignment-based filtering is replaced by shape-based filtering for improved speed. Hits with ZP-scores less than 0.95 are filtered. All retained monomers are then mapped to their respective multimers, yielding a maximum of 1000 multimeric hits. The IF-score for each subunit in the mapped multimers is derived from previously identified similar subunits, set to 0 if not found (indicated by grey cartoon); and the mean IF-score for each mapped multimer is calculated based on all subunits. Finally, multimeric hits from both the IFM and ZPM are consistently ranked using the Q-score as defined in Equation (9), with the top 1000 hits returned at the end of the process.
